# Global dissemination of FC-*ϵ* RI and p53 signalling perturbation contribute to Dasatinib resistance in Pancreatic Cancer Cell-lines

**DOI:** 10.64898/2026.02.22.707324

**Authors:** AKM Azad, Muhammad J. A. Shiddiky, Mohammad Ali Moni

## Abstract

Acquired Dasatinib Resistance (ADR) hinders efficacious treatment of Pancreatic Cancer (PC), often mediated by dynamic signalling reprogramming due to prolonged drug intake. With a novel signalling cross-talk network modelling, this study analyses transcriptomics data of dasatinib-resistant and dasatinib-sensitive pancreatic cancer cell lines and prioritizes key signalling molecules via systemic coordination of their magnitude of dysregulation and the degree of signalling cross-talk among enriched pathways. Results found the *p53* and *FC-ϵ RI signalling* pathways demonstrating significant perturbation enrichment, complementarily orchestrating a total of 87% of the global perturbation map in dasatinib resistance. Further statistical characterization of the cross-talk network identified 10 key resistant biomarkers, including *THBS1, CDKN1A*, and *BCL2L1* within *p53 signalling*, and *RAC2* and *MAPK13* within *FC epsilon RI signalling*. Validation with TCGA transcriptomics, CPTAC relative proteomics, and StringDB protein-protein interaction data for their potential prognostics revealed *BCL2L1* as pivotal for global perturbation dissemination and, thereby, a novel therapeutic target.

## I Introduction

Pancreatic cancer (PC) is one of the most fatal cancers among all others due to its high mortality rate and poor early diagnosis [1]. Despite comprehensive treatment with chemo- and radiotherapy and complete tumour resection, a 5-year survival rate is only 13% [1]. Some chemotherapeutic agents, such as nab-Paclitaxel, gemcitabine, folfirinox, etc., have been used in clinical practices, but increased toxicity and severe-risk imposition upon highly metastatic patients render poor survival outcomes in PC patients [2].

Dasatinib, a multi-kinase inhibitor of Src family kinases commonly used to treat blood and bone marrow cancers, has also been approved by the Food and Drug Administration (FDA) for PC treatment [3] since Src inhibitors have the potential to reduce cell proliferation and induce cell cycle arrest in PC cells [4]. In pancreatic ductal adenocarcinoma (PDAC) cell lines (Panc-1 and Colo-357), dasatinib effectively blocks tumour stem cell generation via suppressing the expression of transforming growth factor-beta (TGF-*β*) target genes that are related to epithelial-mesenchymal transition and invasion [5].

Acquired resistance to targeted therapies emerges during the treatment of many cancers, including PC [6, 7], where the adaptive tumour microenvironment and epigenetic remodelling play crucial roles. In murine PDAC cells, aberrant methylation patterns were observed in multiple genomic sites of regulatory elements under increasing *MEK*_*i*_ treatment (MAPK pathway inhibitor), suggesting the epigenetic remodelling along with a single clonal expansion is responsible for its resistance [6]. Moreover, perturbation in signalling pathways leads to alternate activation of the compensatory signalling network [8, 9]. This phenotypic plasticity facilitates cancer cells to evade apoptosis, sustain growth, and ultimately thrive in the presence of targeted therapy [8, 9].

Precise mechanisms underpinning acquired resistance (AR) to a drug require highly contextual analysis of relevant omics data. Therefore, we hypothesized that novel biomarkers contributing to Acquired Dasatinib Resistance (ADR) could be identified by seeking answers to the following research questions: 1) *which signalling proteins/genes/molecules are dysregulated in dasatinib-resistant vs dasatinib-sensitive cell lines?*, 2) *are their orchestrated activities involving the pathways (enriched with dysregulated entities) via signalling cross-talk?*, 3) *any particular signalling protein/-gene/molecule that is driving the whole signalling perturbation?*. This study presents a comprehensive perturbation map of dasatinib resistance by leveraging high-throughput transcriptomic profiling of pancreatic cancer cell-line models. Initial screening of ADR biomarkers and their functional footprints is done via differential expression analysis, followed by their functional enrichment and signalling pathway impact analyses. Moreover, our approach identifies a novel map of highly aberrant perturbation drivers by employing a novel network-based modelling of signalling cross-talk networks. This has the potential to uncover key hubs and bottlenecks that drive resistance by disseminating the perturbation globally. The findings provide valuable insights into the molecular underpinnings of dasatinib resistance and establish a framework for the rational design of next-generation combination therapies for pancreatic cancer.

## 2 Materials & Methods

Overall schematic diagram of our analysis workflow is shown in Figure 1.

**FIGURE 1.**
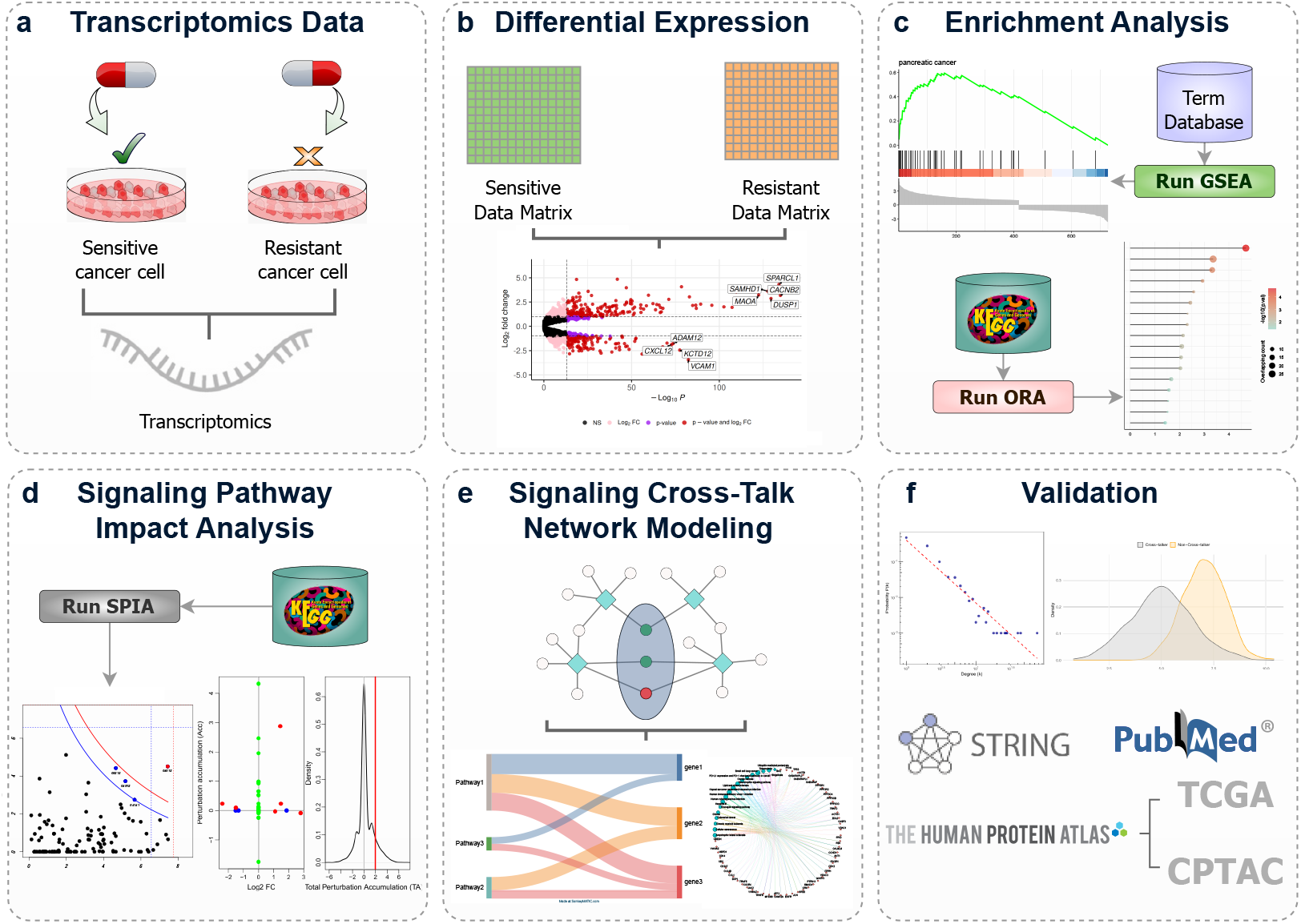
Overall schematic diagram of our proposed workflow in identifying biomarkers of Acquired Dasatinib Resistance (ADR).

### Data Collection & Pre-processing

Gene expression data for Dasatinib-sensitive cell lines (Panc0504, Panc0403, Panc1005) and Dasatinib-resistant cell lines (MiaPaCa2, Panc1, SU8686) are collected from Gene Expression Omnibus (GEO) repositories with the accession number GSE59357. This dataset used three biological replicates to extract total RNA using complementary DNA microarray technology with the Illumina Human HT-12 v4 BeadChip (GPL10558). Two datasets (i.e., resistant and sensitive) were constructed via log2-transformation and quantile-normalization before further analyses were conducted.

### Differential expression analysis

To identify a list of differential biomarkers, linear models were employed to resistant-vs-sensitive contrast matrices with empirical Bayes function and vooma transformation. Statistically significant differential expression changes were selected based on two thresholds: 1) *abs*(*log*2*FC*) ≥ 1.0 (log2 fold change) and 2) FDR-adjusted *P*.*value* ≤ 0.05.

### Gene signature analysis

To observe the predominance of known gene signatures (as a predefined list of genes) in identified differentially expressed genes (Resistant-vs-Sensitive conditions), the GSEA (Gene Set Enrichment Analysis) was performed. It was hypothesized that Pancreatic cancer-related known gene signatures will be highly prevalent with differentially expressed genes in dastatinib resistance. The *gseDO* function from *DOSE* R package was used for this analysis, where the parameter values that were considered are: *nPerm=1000* (permutation iteration), *minGSSize=10* (minimal size of each geneSet), *maxGSSize=500* (maximum size of each geneSet), *pvalueCutoff=0*.*05*, and *pAdjustMethod=“BH”* (benjamini-hochberg).

### Functional pathway over-representation analysis

Enrichment of differentially expressed genes with known signalling pathways indicates strong relevance of their functional coherence. Therefore, a set of 48 signalling pathways from the KEGG (Kyoto Encyclopedia of Genes and Genomes) database [10] were collected for this purpose. To identify which signalling pathways are highly enriched with identified DEGs, a *Hypergeometric test*-based over-representation analysis is conducted using *phyper* function (with *lower*.*tail* = *FALSE*) from *stats* R package with all the genes within the study as the background geneset. Statistically significant signalling pathways are defined with the enrichment *P*.*value* ≤ 0.05 from the Hypergeometric test using the Equation **??**. The geneRatio values of each enriched pathway were computed by determining the ratio between the number of enriched DEGs and the size of the corresponding pathway.

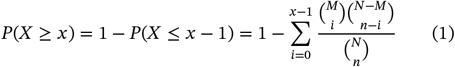

where *N* = the total number of genes in the background gene list, *M* = genes in the Pathway, *n* = the number of DEGs used for this analysis, and the observed number *x* = the number of DEGs found within the given pathway. Here, the generated *P*.*value* is a right-tail probability, i.e., *Pr*(*X* ≥ *x*), which determines the chance of having a larger number of overlaps than *x*. Hence, a smaller *P*.*value* would indicate that the observed number of DEGs in the pathway is significantly higher than expected by chance, suggesting enrichment of the pathway with DEGs.

### Cross-talk identification among enriched signalling pathways

An inter-pathway cross-talk network was constructed to examine systemic interactions among enriched signalling pathways via shared genes. Let *G* := (*V, G*) be a signalling cross-talk network model, where *V* is the set of nodes representing either DEGs or pathways and *E* is the set of edges denoting the gene-pathway relationships obtained from signalling transduction pathways from KEGG database. Network construction was performed using the *igraph* R package. Cross-talking DEGs were defined as those connecting at least two pathways. Next, to identify perturbed and cross-talking DEGs, the network was filtered to retain only significant nodes (DEGs with *abs*(*log*_2_*FC*) ≥ 2) and pruned to exclude low-degree nodes (degree = 1). Node degree distribution was computed to observe the scale-free properties of the network. Central hubs, namely the *resistant biomarkers* were identified as nodes with high connectivity (the number of pathways cross-talking via it) and dysregulation (*abs*(*log*_2_*FC*)). Pearson’s correlation was measured to quantify correlation between degree of the *resistant biomarkers* and their *log*_2_*FC*. A *two-sided* and *two-sample Kolmogorov-Smirnov test* was conducted (using *ks*.*test* function from *base* R package) to test if the magnitude of differential expression of *resistant biomarkers* (i.e., cross-talking hubs) and its counterparts are drawn from the same distribution. The biological significance of hubs was assessed through literature mining and experimental evidence.

### Signalling Pathway Impact Analysis

Over-representation-based enrichment analysis is a topology-unaware approach, which may lack mechanistic consideration. Therefore, the Signaling Pathway Impact Analysis (SPIA) was performed to quantify pathway perturbations integrating both over-representation of DEGs and pathway topology-driven signalling perturbation. It is hypothesized that the pathway perturbation through constituent signalling proteins depends both on their differential expressions and the perturbation activities of their upstream signalling proteins [11]. SPIA computes two probabilities: (1) *pN*_*DE*_, representing the over-representation of DEGs in a pathway using a hypergeometric test [see Equation **??**], and (2) *pPERT*, quantifying the pathway perturbation propagated by DEGs through associated signalling links (activation or repression). The null hypothesis for *pPERT* is that the total net accumulated perturbation *tA* [Equation 2] of a particular pathway is more than that of the observed one by chance [Equation 3], where the *T*_*A*_ is total net accumulated perturbation score of a particular bootstrap iteration, where the number of DEGs remains the same as an observed case, but to topology and differential expression fold-changes both were randomized.

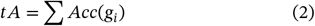

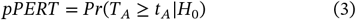

where the *Acc*(*g*_*i*_ ) is the net perturbation of a gene, *g*_*i*_, calculated by the following Equation: [Equation 4]

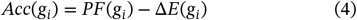

where, Δ*E*(*g*_*i*_) is the normalized expression changes of the gene *g*_*i*_ in resistant-vs-sensitive condition. Next, the *PF*(*g*_*i*_) is the perturbation factor of the gene *g*_*i*_, which is determined by the following Equation [Equation 5]:

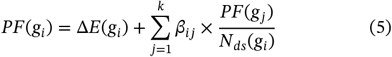

where *N*_*ds*_(*g*_*j*_) is the number of downstream genes of *g*_*j*_, and *β*_*ij*_ is a coefficient indicating the type of signalling link, i.e., activation or inhibition.

Next, for each pathway perturbation significance, a global p-value, namely *pG*, is determined by combining *pNDE* and *pPERT* by the “normal inversion” technique by setting the *combine* parameter as ‘*norminv*’ [Equation 6].

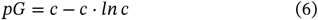

where *c* = *pNDE* ⋅ *pPERT*. The global p-value *pG*, is further corrected using both FDR and Bonferroni correction methods, termed as *pGFDR* and *pGFWER*, respectively.

### Visual interpretation of Pathway perturbation using PathView

To understand the perturbation effects of DEGs in a given enriched pathway, the *pathview* function from the *PathView* R package is used. PathView is a Bioconductor-hosted pathway analysis toolset which facilitates the seamless integration of differential expression results (user-supplied) and overlays the dysregulation information on the known pathway annotations. It automatically downloads the annotated genes, molecules and topology (inter-connections), parses it and renders the overlaid differential expression information.

### Validation with Existing knowledge

Biomarkers are evaluated with multiple information sources - protein-protein association information from StringDB [12]. Evidence of potential prognostic values from gene expression changes from TCGA mRNA expression changes in Human Protein Atlas database [13]. StringDB web portal facilitates queries with multiple genes of a chosen species to output an induced sub-network with potential protein-protein interactions among them, defined by various sources of evidence, e.g., known interactions (database curated and experimentally verified), predicted (gene neighbourhood, gene fusions, and gene co-occurrence), text-mining, co-expression, and protein homology [12]. It also offers a set of network statistics regarding the output, including node and edge counts, average node counts, average local clustering coefficient (LCC), and statistical significance of the induced sub-network against the background (by randomizing the gene sets with the same size and degree distributions). Moreover, to facilitate further clustering, more nodes with their incident interactions from the whole PPI network (involving other genes in the whole genome) are augmented. For clustering the stochastic flow-based MCL (Markov Clustering) algorithm is employed with the inflation parameter thresholding. For reinforcement of the biological relevance of identified resistant biomarkers, further enrichment was conducted using GO terms (biological process), Reactome pathways, WikiPathways, Subcellular localizations information, and their Pubmed queries.

The Human Protein Atlas database has the gene expression from TCGA (The Cancer Genome Atlas) and protein expression from CPTAC (Clinical Proteomic Tumor Analysis Consortium) repositories for pancreatic adenocarcinoma (PAAD). Preprocessed transcriptomics (nRPX) data of Pancreatic cancer from 176 patients (84 alive and 92 deceased) with various stages (I, II, III, and IV) is observed for each biomarker in this study from the HPA database. TCGA RNA expression data from the HPA database were obtained for each of those resistant biomarkers. Using a “quartile”-based dichotomized of each gene expression (i.e., High or Low expression), survival analysis is performed using *survminer* R package. Tandem mass tag (TMT) mass spectrometry-based CPTAC proteomics data from 145 tumours and 90 normal was preprocessed, and relative proteomics expression was reported [13]. Relative protein expression of biomarkers genes from CPTAC data in Tumor-vs-Normal conditions were compared using *Wilcoxon test* to observe their differential abundance using *ggbetweenstats* R function (*ggstatsplot* R package [14]).

## 3 Results

Dasatinib is a dual inhibitor of Abl/Src tyrosine kinases, normally used for treating Ph-positive leukaemia, but has shown competence in Pancreatic ductal adenocarcinoma (PDAC) in preclinical studies [15]. However, its efficacy in clinical settings wasn’t promising, primarily due to the acquired resistance [16].

In this study, we have investigated the role of differential biomarkers in signalling perturbation in resistance-vs-sensitive cell lines and their impact on dasatinib resistance. We hypothesized that key signalling biomarkers and their network-based signalling impact analysis might reveal the putative mechanisms of acquired resistance to dasatinib in Pancreatic cancer cell lines.

### Differential Expression Analysis and GSEA-based Initial Insights into Biological Relevance

We collected gene expression datasets of pancreatic cancer cell lines with Dastatinib-sensitive and Dastatinib-resistant conditions, which were further preprocessed using log2-transformation and quantile-normalization using *limma* R package. Then, we performed differential expression analysis in resistant-vs-sensitive using linear models with empirical Bayes with *vooma* using *limma* R package. After applying a combined threshold of *abs*(*log*_2_*FC*) ≥ 1.0 (log2 fold change) and FDR-adjusted *P*.*value* ≤ 0.05, we have obtained a total of 727 differentially expressed genes (DEGs) [Supplementary table 1], among which 417 genes were up-regulated and 310 were down-regulated. Top 20 DEGs (based on *log*_2_*FC*) are shown in Table 1, where VIM, CBS, SRGN, TUBB2B, and TM4SF18 are some highly downregulated DEGs, and S100P, TACSTD2, LCN2, KRT7, and MSLN are a few highly upregulated DEGs [Figure 2a]. To observe the biological relevance of the identified DEGs in acquired dasatinib resistance in pancreatic cancer, we have conducted a GSEA (Gene Set Enrichment Analysis) using *fgsea* with minimum and maximum size of each geneset 10 and 500, respectively. Highly enriched biological terms were selected based on a filter: Benjamini-Hochberg-adjusted adjusted *P*.*value* < 0.0001, yielding 14 terms. Among these, the most prominently enriched terms include: “*pancreatic cancer*” [normalized enrichment score (NES) = 3.03, FDR = 6.4 × 10^−10^, 27 DEGs matches], “*pancreatic carcinoma*” [NES = 3.01, FDR = 1.7 × 10^−8^, 22 DEGs matches] and “*pancreatic adenocarcinoma*” [NES = 2.75, FDR = 4.02×10^−7^, 15 DEGs matches] [Figure 2b], where the visualization of distributions of running enrichment scores with pre-ranked gene list (727 DEGs) suggests that top terms are shown highly enriched with upregulated genes [Figure 2c]. These results suggest that significant DEGs are highly contextual towards pancreatic cancer metastasis and/or acquired resistance towards dasatinib drug in pancreatic cancer.

**TABLE 1.**
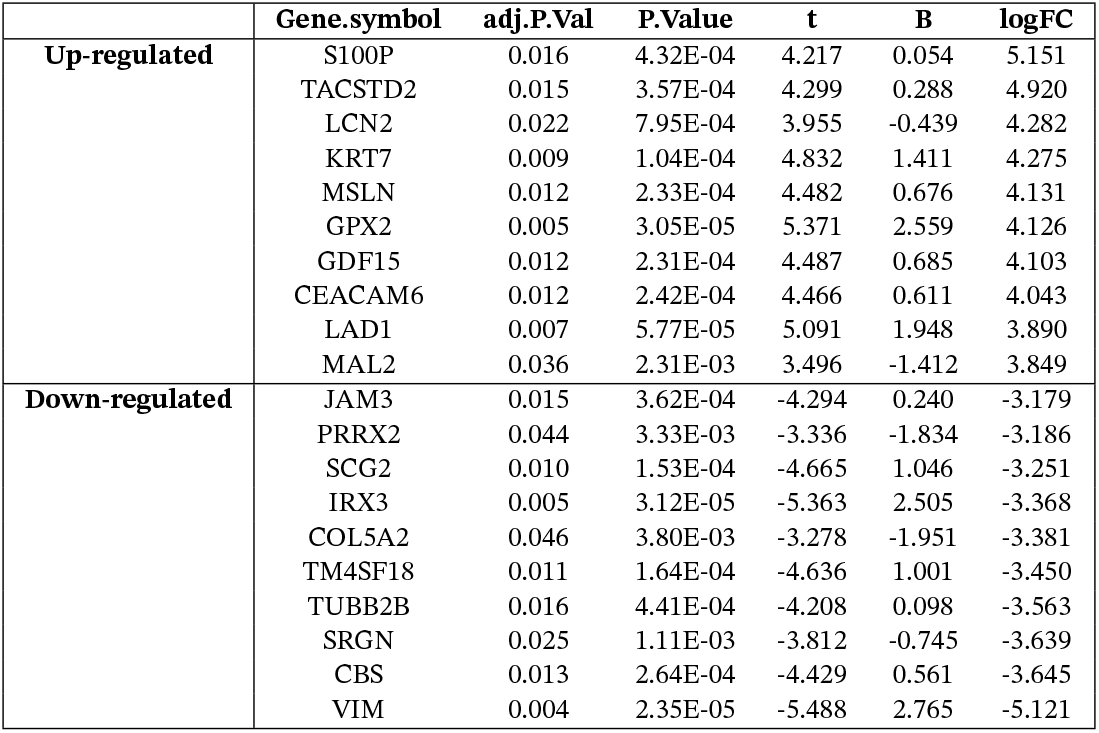
Top 20 up-regulated and down-regulated DEGs in Acquired Dasatinib resistance.

**FIGURE 2.**
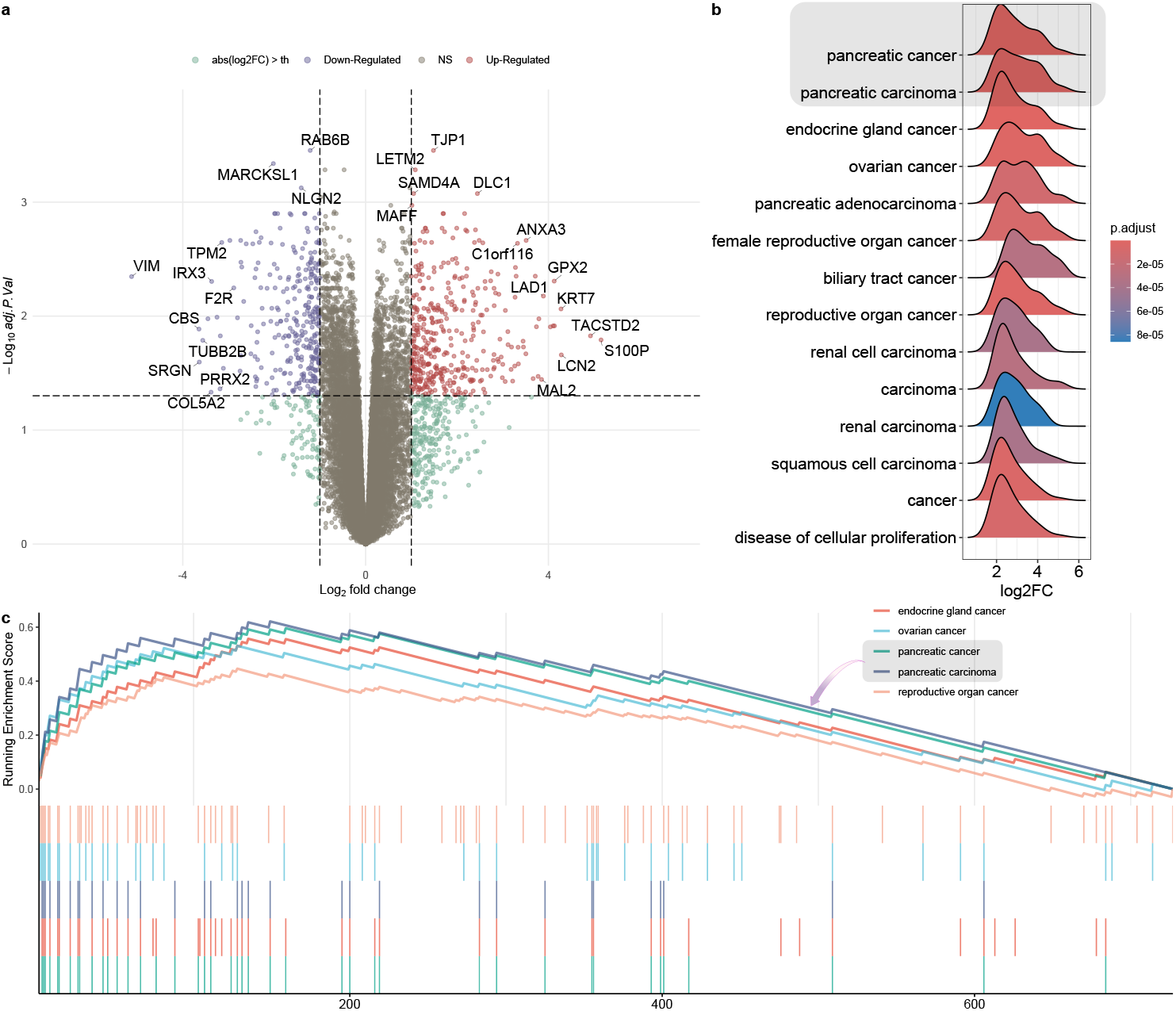
Differential Gene Expression Analysis. **(a)** Volcano plot shows genes are differently expressed in acquired Dasatinib resistance-vs-sensitive conditions. **(b)** Geneset Enrichment analysis (GSEA) of 727 selected DEGs (*abs*(*log*_2_*FC*) ≥ 1.0 and FDR-adjusted *P*.*value* < 0.05), where most of the top enriched terms were pancreatic cancer-related, revealing the biological relevance of such differential expression in resistance-vs-sensitive conditions. **(c)** The distributions of running enrichment scores with pre-ranked gene list (727 DEGs) demonstrate that upregulation of pancreatic cancer as the gene-hits are clustered toward positive ranking metric, which indicates gene upregulations.

### Over-Representation Analysis of KEGG Signaling Pathways

To determine the signalling perturbation effects of the DEGs, we performed an over-representation-based pathway enrichment test (i.e., Hypergeometric test) against known 48 KEGG signalling pathways, where the whole list of genes in this cohort was considered as the background gene set. Using a threshold of enrichment *P*.*value* ≤ 0.05 (from Hypergeometric test), we obtained 17 signalling pathways as significantly enriched with the DEGs [Fig 3a]. Among those, *p53, Fc epsilon RI, VEGF, PPAR, NF-κB*, and *MAPK* signalling pathways were highly enriched based on the enrichment gene ratio [Fig 3b], which is a proportion of pathway genes that are enriched in a given pathway [Table 2].

**TABLE 2.** DEG-enriched KEGG Signalling pathways. Pathways are ordered based on overlapping GeneRatio (proportion of pathway genes that are enriched in a given pathway)

| Description | Count | GeneRatio | pvalue | p.adjust |
| --- | --- | --- | --- | --- |
| p53 signaling | 8 | 0.107 | 0.001 | 0.013 |
| Fc epsilon RI signaling | 7 | 0.101 | 0.003 | 0.024 |
| VEGF signaling | 6 | 0.100 | 0.005 | 0.027 |
| PPAR signaling | 7 | 0.092 | 0.005 | 0.027 |
| NF-kappa B signaling | 9 | 0.086 | 0.004 | 0.027 |
| MAPK signaling | 25 | 0.083 | 0.000 | 0.001 |
| Phosphatidylinositol signaling system | 8 | 0.082 | 0.007 | 0.034 |
| Rap1 signaling | 17 | 0.080 | 0.000 | 0.007 |
| TNF signaling | 9 | 0.076 | 0.009 | 0.034 |
| Neurotrophin signaling | 9 | 0.075 | 0.009 | 0.034 |
| B cell receptor signaling | 6 | 0.071 | 0.029 | 0.084 |
| ErbB signaling | 6 | 0.070 | 0.030 | 0.084 |
| PI3K-Akt signaling | 25 | 0.069 | 0.000 | 0.007 |
| Wnt signaling | 12 | 0.069 | 0.008 | 0.034 |
| Sphingolipid signaling | 8 | 0.066 | 0.027 | 0.084 |
| Hippo signaling | 10 | 0.064 | 0.022 | 0.074 |
| Relaxin signaling | 8 | 0.062 | 0.038 | 0.099 |

**FIGURE 3.**
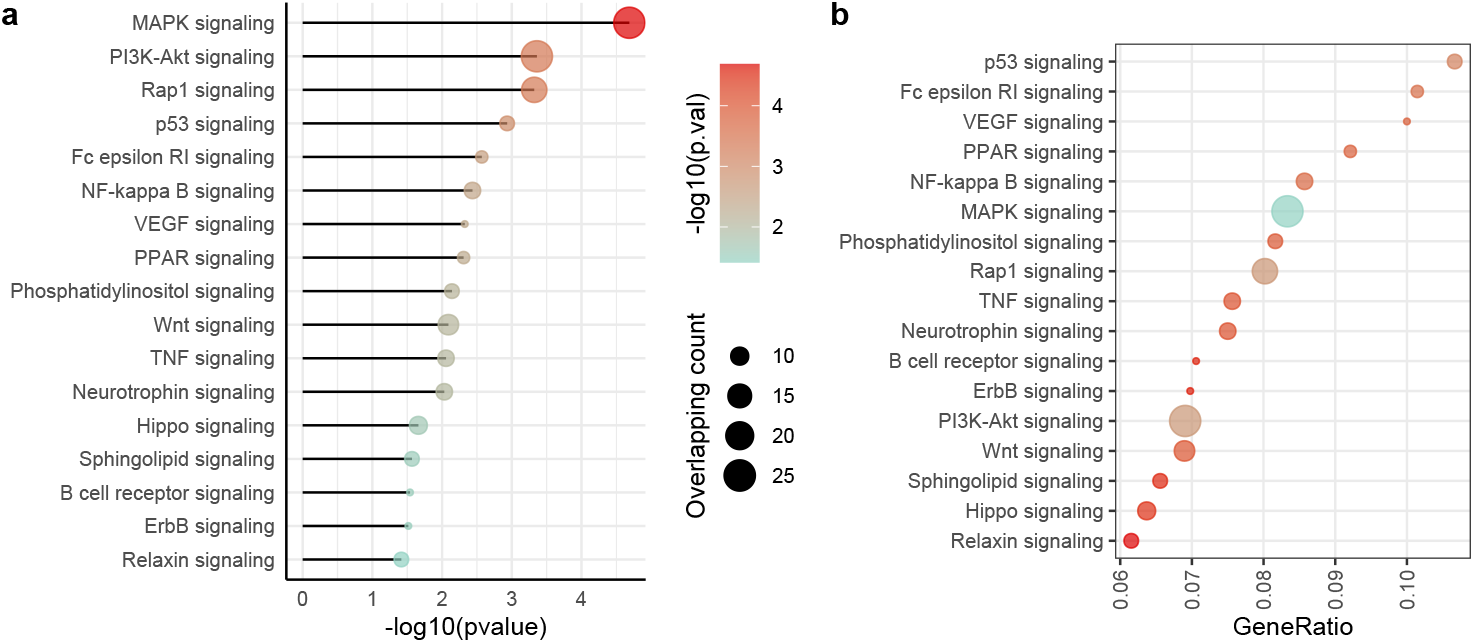
Functional enrichment analysis of DEGs with known signalling pathways from KEGG. DEGs (adjusted p-value < 0.01) were subjected to the signalling pathway (KEGG) enrichment analysis using a Hypergeometric test - revealing significantly (*P*.*value* ≤ 0.05) enriched signalling pathways with high enrichment geneRatio values, including p53, Fc epsilon RI, VEGF, PPAR, and NF-kappa B signalling pathways.

### P53 and FC epsilon RI Signaling Cross-Talks are crucial in disseminating overall perturbation

Pathway enrichment tests reveal important functional footprints of differential biomarkers. However, it lacks the insights of detailed coordination among those enriched functional terms. Therefore, we performed a signalling cross-talk analysis among 17 top enriched KEGG signalling pathways, where the cross-talk between two pathways was identified via a shared component (gene/protein/molecule) that is a constituent of both [17, 8, 9, 18]. Figure 4a shows the network view of these signalling crosstalks, how the involved pathways are interconnected and share their perturbations. In this network, the nodes are either DEGs and their enriched signalling pathways, and the edges are the pathway-DEG memberships. Hence, any DEG that is shared between two pathways is considered to be a cross-talking bridge between those pathways. Note that this network is constructed in an ad-hoc manner, which doesn’t necessarily indicate physical protein-protein interaction, and regarding the DEGs in this network, the ones that are not shared with at least two pathways are pruned out. Thus, we obtained 16 signalling pathway nodes and 34 DEG nodes in this network [Figure 4a]. Here, pathway node sizes are proportionate to the number of DEGs involved, and DEG node colours are proportionate to the degree of their log2 fold changes. We were particularly interested in observing the signalling pathways that were found to be highly enriched with higher geneRatio (see Table 2), as it highlights the pathways in a size-invariant manner. As *p53* and *FC epsilon RI signalling* pathways were shown higher geneRatio than others (post-filtration with enrichment *P*.*value* < 0.05), we investigated those two pathways further. Table 3 and Table 4 show *p53* and *FC epsilon RI signalling* pathways and their cross-talking pathways, respectively, via their shared DEGs and their corresponding *log*_2_-fold changes and *adjusted*-*P*.*values*. It reveals that *p53 signalling* pathway was involved with a total of 6 DEGs (**CDKN1A, CCND1, GADD45A, PPP1R13L, BCL2L1**, and **THBS1**), involving 7 of 16 signalling pathways (43.8%) [Figure 4b]. Regarding *FC epsilon RI signalling* pathways, a total of 7 DEGs (**LYN, INPP5D, RAC1, RAC2, MAP2K1, MAPK13**, and **FYN**) are defined as cross-talks involving 13 of 16 signalling pathways (81.2%) [Figure 4b]. Interestingly, except the “*Hippo signalling*”, 6 others that were found to be cross-talking with *p53 signalling* pathway were also found to be within 13 cross-talking pathways with *FC epsilon RI signalling*. This indicates that *p53* and *FC epsilon RI signalling* pathways combinedly cross-talk with 14 out of all 16 signalling pathways in this network, which is 87.5% [Figure 4b].

**TABLE 3.** p53 signalling pathways cross-talking with other signalling pathways via highly up-regulated shared genes.

| Pathway1 | Pathway2 | Shared Gene | log2FC | adj.P.Val |
| --- | --- | --- | --- | --- |
| p53 signaling | ErbB signaling | CDKN1A | 2.738 | 0.034 |
|  | Hippo signaling | CCND1 | 1.111 | 0.013 |
|  | MAPK signaling | GADD45A | 1.280 | 0.011 |
|  | NF-kappa B signaling | PPP1R13L | 1.336 | 0.018 |
|  | NF-kappa B signaling | GADD45A | 1.280 | 0.011 |
|  | NF-kappa B signaling | BCL2L1 | 1.993 | 0.002 |
|  | PI3K-Akt signaling | THBS1 | 3.336 | 0.019 |
|  | PI3K-Akt signaling | CCND1 | 1.111 | 0.013 |
|  | PI3K-Akt signaling | CDKN1A | 2.738 | 0.034 |
|  | PI3K-Akt signaling | BCL2L1 | 1.993 | 0.002 |
|  | Rap1 signaling | THBS1 | 3.336 | 0.019 |
|  | Wnt signaling | CCND1 | 1.111 | 0.013 |

**TABLE 4.** FC epsilon RI signalling pathways cross-talking with other signalling pathways via DEGs.

| Pathway1 | Pathway2 | Shared Gene | log2FC | adj.P.Val |
| --- | --- | --- | --- | --- |
| Fc epsilon RI signaling | B cell receptor signaling | LYN | -1.061 | 0.019 |
|  | B cell receptor signaling | INPP5D | 1.266 | 0.015 |
|  | B cell receptor signaling | RAC1 | 1.491 | 0.038 |
|  | B cell receptor signaling | RAC2 | 3.506 | 0.021 |
|  | B cell receptor signaling | MAP2K1 | 1.106 | 0.019 |
|  | ErbB signaling | MAP2K1 | 1.106 | 0.019 |
|  | MAPK signaling | MAP2K1 | 1.106 | 0.019 |
|  | MAPK signaling | RAC1 | 1.491 | 0.038 |
|  | MAPK signaling | RAC2 | 3.506 | 0.021 |
|  | MAPK signaling | MAPK13 | 2.801 | 0.029 |
|  | NF-kappa B signaling | LYN | -1.061 | 0.019 |
|  | Neurotrophin signaling | RAC1 | 1.491 | 0.038 |
|  | Neurotrophin signaling | MAPK13 | 2.801 | 0.029 |
|  | Neurotrophin signaling | MAP2K1 | 1.106 | 0.019 |
|  | PI3K-Akt signaling | MAP2K1 | 1.106 | 0.019 |
|  | PI3K-Akt signaling | RAC1 | 1.491 | 0.038 |
|  | Phosphatidylinositol signaling | INPP5D | 1.266 | 0.015 |
|  | Rap1 signaling | RAC1 | 1.491 | 0.038 |
|  | Rap1 signaling | RAC2 | 3.506 | 0.021 |
|  | Rap1 signaling | MAP2K1 | 1.106 | 0.019 |
|  | Rap1 signaling | MAPK13 | 2.801 | 0.029 |
|  | Relaxin signaling | MAPK13 | 2.801 | 0.029 |
|  | Relaxin signaling | MAP2K1 | 1.106 | 0.019 |
|  | Sphingolipid signaling | RAC1 | 1.491 | 0.038 |
|  | Sphingolipid signaling | RAC2 | 3.506 | 0.021 |
|  | Sphingolipid signaling | MAPK13 | 2.801 | 0.029 |
|  | Sphingolipid signaling | FYN | 1.410 | 0.021 |
|  | Sphingolipid signaling | MAP2K1 | 1.106 | 0.019 |
|  | TNF signaling | MAPK13 | 2.801 | 0.029 |
|  | TNF signaling | MAP2K1 | 1.106 | 0.019 |
|  | VEGF signaling | RAC1 | 1.491 | 0.038 |
|  | VEGF signaling | RAC2 | 3.506 | 0.021 |
|  | VEGF signaling | MAPK13 | 2.801 | 0.029 |
|  | VEGF signaling | MAP2K1 | 1.106 | 0.019 |
|  | Wnt signaling | RAC1 | 1.491 | 0.038 |
|  | Wnt signaling | RAC2 | 3.506 | 0.021 |

**FIGURE 4.**
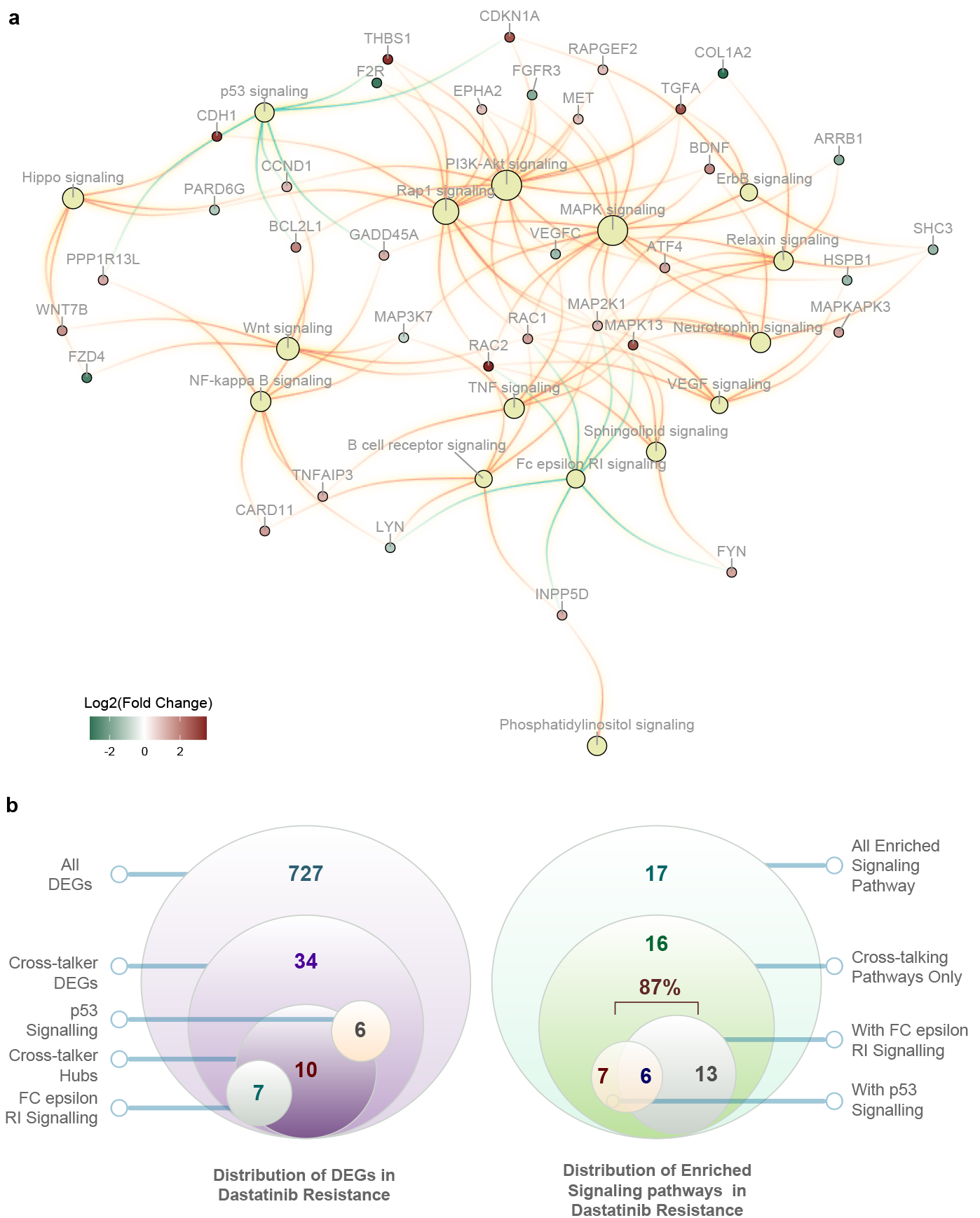
Signalling cross-talks among DEG-enriched KEGG signalling pathways. **(a)** Enriched signalling pathways were found to be cross-talking among themselves involving highly over-expressed (e.g., SHC3, TGFA, THBS1, BDNF, BCL2L1, WNT7B, CCND1, and VEGFC ) genes and under-expressed (e.g., PPP1R13L, RAC1, RAC2, CARD11, F2R, FYN, LYN, and FZD4 ) in Dasatinib-sensitive versus Dastatinib-resistant conditions. p53 and FC epsilon RI signalling pathways (green colour edges) - two of the most highly enriched pathways (based on geneRatio values) were highly complementary in coordinating these cross-talks as they shared no constituent genes between them. Note, each genes are also coloured in proportion to their differential expression level (i.e., *log*_2_-scale) **(b)** Distribution of cross-talker DEGs and their enriched Signalling pathways, particularly in p53 and FC epsilon signalling pathways which combinedly cross-talks with 87% of all signalling pathways involved, indicating the dominance in driving global perturbation.

### Scale-Free Topology Identifies Key Crosstalk Hubs Driving Resistance-Induced Dysregulation Cascade

Next, we investigated the biological relevance and analytical rigour of such gene-pathway cross-talk [see 3] by observing the potential existence of its scale-free topology. Therefore, we determined the degrees of all the network nodes and observed their frequency distribution. The results suggest that a few nodes (hubs) exhibit disproportionately high connectivity compared to the rest [Figure 5a], a hallmark of scale-free topology in biological networks [19]. Moreover, we observed similar behaviour when we only selected DEG nodes (ignoring their associated pathway nodes) [Figure 5b].

**FIGURE 5.**
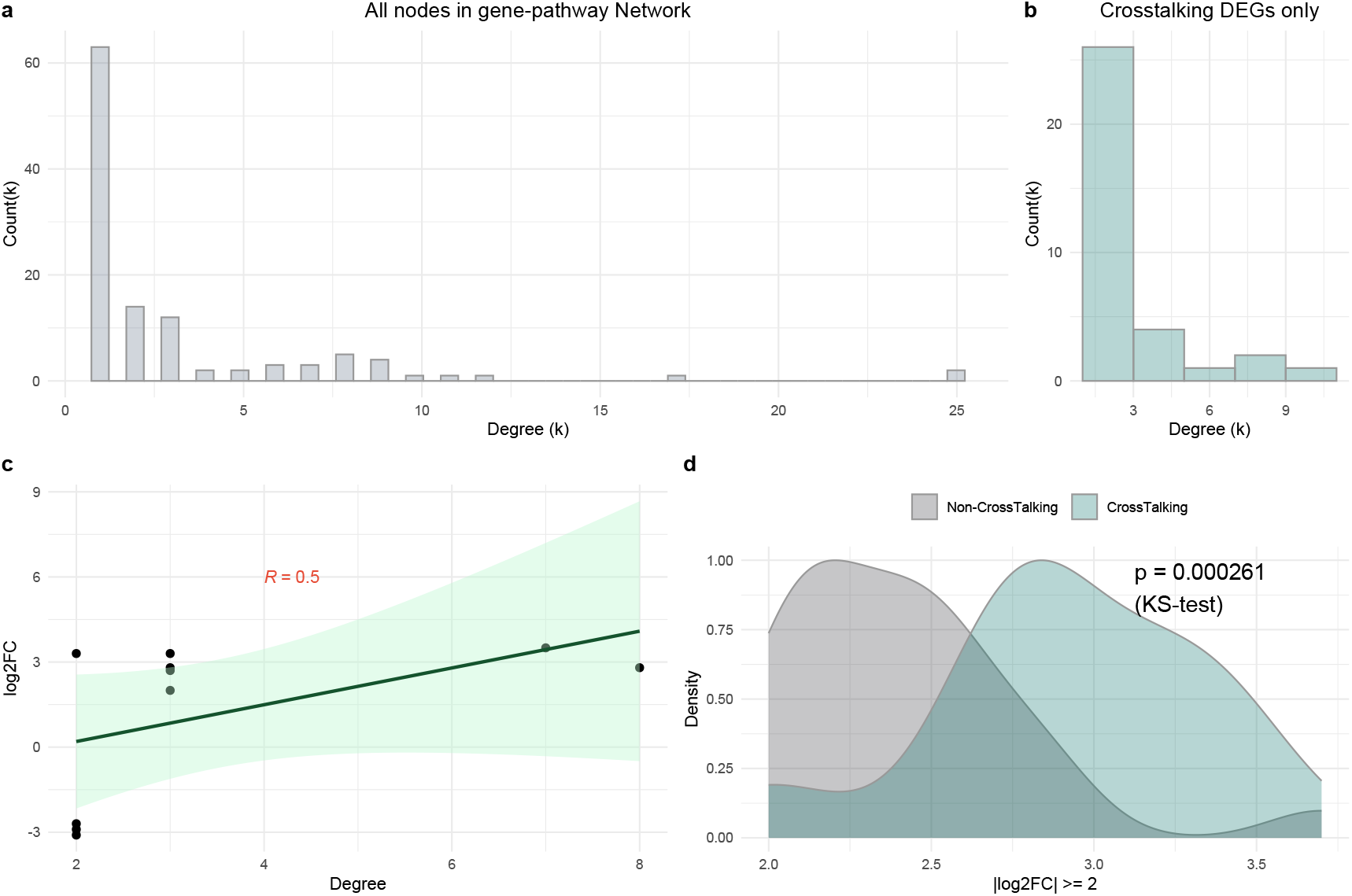
Scale-Free topology of cross-talk network and the association of its cross-talking hubs with their differential expression in resistance conditions. **(a)** Gene-Pathway cross-talk network reveals the scale-free topology where only a few nodes have very large connectivity. **(b)** Similar scale-free pattern was observed when only cross-talking DEGs (*degree* ≥ 2) were considered, **(c)** For the cross-talking DEGs (*degree* ≥ 2) that have high differential expression (*log*_2_*FC* ≥ 2 or *log*_2_*FC* ≤ −2) (a total of 9 DEGs), a moderate correlation (*R* = 0.51) was observed between their degree of connectivity and degree of dysregulation. **(d)** Moreover, compared to the non-cross-talking DEGs, those 9 cross-talking DEGs demonstrated a statistically significant difference in the distribution of dysregulation when a filter of *abs*(*log*_2_*FC*) > 2) was applied for all the DEGs in the network.

Having the scale-free property within the network topology, it is also worth investigating the association of differential expression changes of DEGs in resistant conditions compared to sensitive conditions. This is crucial for prioritizing hub cross-talks (i.e., DEGs) with high-degree centralities that exhibit significant differential expression as those may not only be key regulators of resistance-causing perturbations but also be targeted by therapeutics. In that regard, we investigated the cross-talking DEGs (i.e., *degree* ≥ 2, shared by at least two pathways) that have higher differential expression in resistant-vs-sensitive cell lines (i.e., *log*_2_*FC* ≥ 2 or *log*_2_*FC* ≤ −2) than others. This selection strategy yielded a total of 10 cross-talking DEGs, which revealed a moderate correlation (*R* = 0.5) between the node degrees of cross-talking DEGs (*degree* ≥ 2) and their corresponding *abs*(*log*_2_*FC*) values [Fig 5c]. Those include **COL1A2, F2R, FZD4, BCL2L1, CDKN1A, TGFA, MAPK13, THBS1, CDH1**, and **RAC2** [Table 5]. We hypothesize these 10 cross-talking hubs as notable biomarkers in dasatinib resistance, where the first 3 genes are down-regulated, and the remaining are up-regulated in resistant-vs-sensitive conditions, respectively. Moreover, a non-parametric test, i.e., *KolmogorovSmirnov test* revealed that the distribution of *abs*(*log*_2_*FC*) values (in resistant-vs-sensitive conditions) for those 9 cross-talking DEGs (*degree* ≥ 2) is significantly differ (*P*-value = 0.000261) from those of non-cross-talking DEGs (*degree* = 1). This may further reinforce that cross-talking, and Hub DEGs are not only perturbed by themselves but also their dysregulation has a cascading effect, perturbing multiple pathways simultaneously. This may significantly contribute to the mechanisms of dasatinib resistance in pancreatic cancer and, therefore, can be defined as highly prioritized biomarkers and novel drug targets.

**TABLE 5.** 10 Cross-talking hubs as significant biomarkers in Dasatinib resistance.

| Biomarkers | Cross-talk degree | Differential fold change |
| --- | --- | --- |
| COL1A2 | 2 | -3.1 |
| F2R | 2 | -2.9 |
| FZD4 | 2 | -2.7 |
| BCL2L1 | 3 | 2 |
| CDKN1A | 3 | 2.7 |
| TGFA | 3 | 2.8 |
| MAPK13 | 8 | 2.8 |
| THBS1 | 3 | 3.3 |
| CDH1 | 2 | 3.3 |
| RAC2 | 7 | 3.5 |

### Signaling Pathway Impact analysis of DEGs demonstrates Pathways with Significant Perturbation Activities

Beyond a mere enrichment test of differentially expressed genes through statistical overlap measurement (i.e., over-representation test), pathway topology analysis may consider the position of genes within a pathway that can identify key signalling genes based on their network centrality and connectivity. To detect the perturbation activities of signalling pathways based on the frequency of DEGs and their structural dependencies through static protein-protein interactions, we conducted the signalling pathway impact analysis (SPIA) [11]. It observes how a particular pathway reveals signalling perturbation via capturing probabilities of two types of evidence, i.e., the probability of over-representation of DEGs (*pNDE*) (measures the chance of biological irrelevance) and the probability of abnormal perturbation through network topology (*pPERT*) (measures the amount of perturbation in each pathway) against the background. The signaling pathways from the KEGG database were only used to construct an SPIA object which was later used for analysing significant DEGs along with their log2 fold changes (from our previous result). We have used *normal inversion* approach to combine both *p*−values (*pPERT* and *pNDE*) since it generates the global *p*-value (i.e., *pG*) by having a higher emphasis on both *pPERT* and *pNDE* being as low as possible (i.e., significant). Note the number of bootstrap iterations, *nB*, was used as default [i.e., *nB* = 2000]. This *p*-value is further corrected using both FDR and Bonferroni correction methods as *pGFDR* and *pGFWER*, respectively. Based on a threshold of *pGFWER* < 0.05, 9 signalling pathways were found to be significantly perturbed in dasatinib resistance [Table 6], including p53, Neurotrophin, Fc epsilon RI, PPAR, WNT, NF-kappa B, MAPK, Rap1, PI3K-Akt signalling pathways. Among these 9 singling pathways, only the *p*53 signalling pathway (KEGG Pathway ID: 04115) demonstrated the most pronounced perturbation (*pPERT* = 0.003) [Figure 6a-c]. Up-regulation of THBS1, CDKN1A, CDKN2A, SFN, BCL2L1, GADD45A, PPP1R13L, CCND1, SERPINB5, and IGFBP3, and down-regulation of CASP8 play crucial downstream roles in controlling/mediating cell cycle arrest, apoptosis, metastasis, and angiogenesis inhibition, DNA damage response, and IGF pathway interactions [Figure 7].

**TABLE 6.** SPIA (Signaling Pathway Impact Analysis) for DEGs to identify pathway perturbation activities. Results are shown only with the significantly perturbed pathways (pGFWER < 0.05) in a sorted manner based on the perturbation p-value, pPERT.

| Pathway Name | pNDE | tA | pPERT | pG | pGFdr | pGFWER | Status |
| --- | --- | --- | --- | --- | --- | --- | --- |
| p53 signaling | 2.0E-05 | -11.056 | 0.003 | 6.4E-07 | 1.5E-05 | 3.1E-05 | Inhibited |
| Neurotrophin signaling | 9.8E-05 | 16.754 | 0.109 | 2.3E-04 | 2.2E-03 | 1.1E-02 | Activated |
| Fc epsilon RI signaling | 5.7E-05 | 14.815 | 0.184 | 3.8E-04 | 2.9E-03 | 1.8E-02 | Activated |
| PPAR signaling | 1.3E-04 | 1.974 | 0.217 | 8.7E-04 | 4.6E-03 | 4.2E-02 | Activated |
| Wnt signaling | 1.5E-05 | 7.124 | 0.294 | 4.3E-04 | 2.9E-03 | 2.1E-02 | Activated |
| NF-kappa B signaling | 2.9E-05 | -7.492 | 0.311 | 7.1E-04 | 4.3E-03 | 3.4E-02 | Inhibited |
| MAPK signaling | 8.6E-12 | 8.033 | 0.363 | 2.8E-07 | 1.3E-05 | 1.3E-05 | Activated |
| Rap1 signaling | 2.6E-08 | 7.620 | 0.627 | 1.5E-04 | 1.8E-03 | 7.0E-03 | Activated |
| PI3K-Akt signaling | 4.4E-10 | 8.351 | 0.642 | 2.3E-05 | 3.6E-04 | 1.1E-03 | Activated |

**FIGURE 6.**
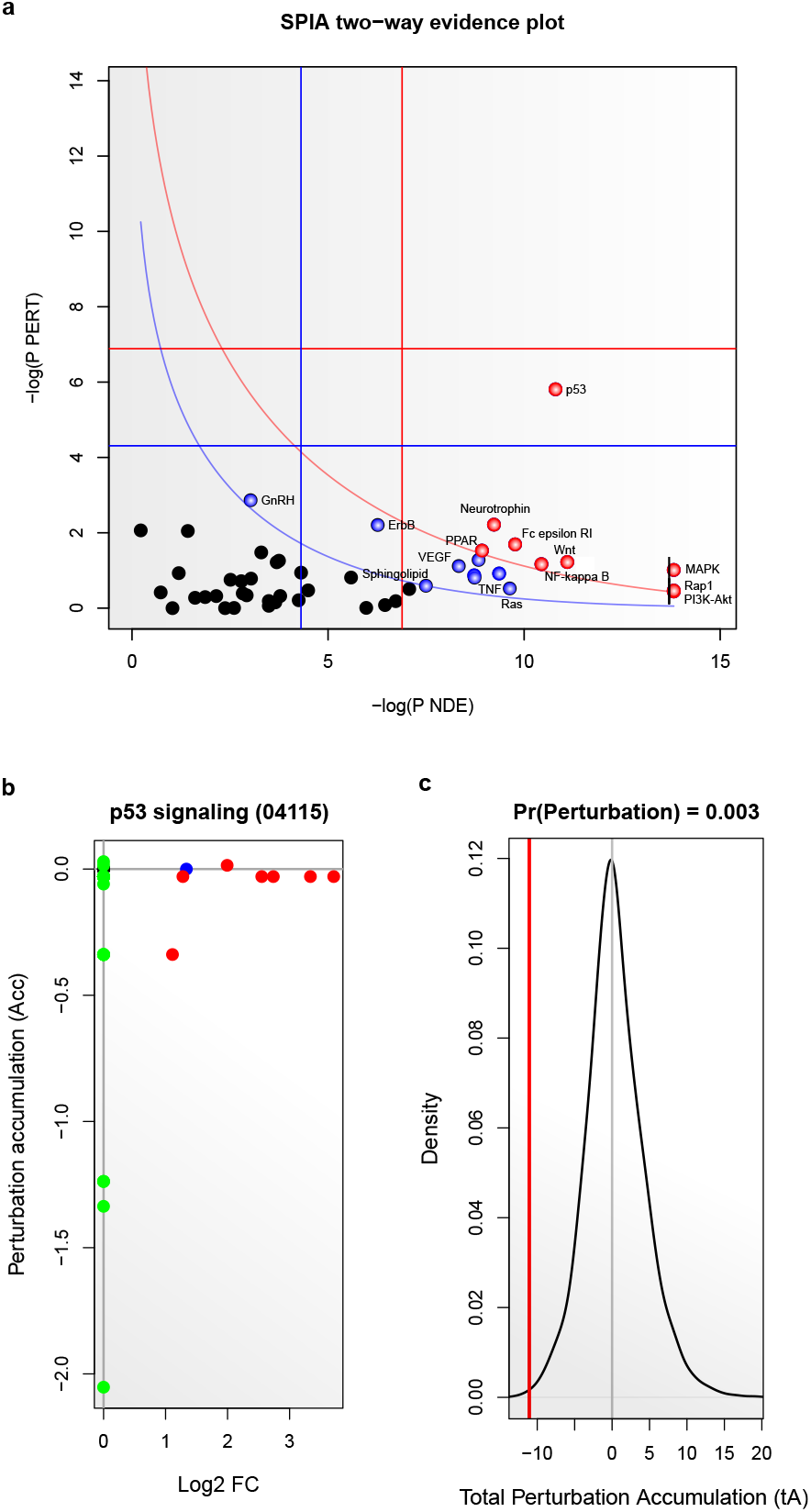
Signaling Pathway Impact Analysis (SPIA) for using the DEGs (i.e., log fold changes) on 49 KEGG signalling pathways topologies. **(a)** By combining the over-representation probability (pNDE) with DEGs and topology-based Perturbation probability (pPERT), the two-way evidence plot reveals the ErbB, Estrogen, PI3K-Akt, and GnRH signalling pathways (red circles) as statistically significant (Bonferroni-corrected *P*-value < 0.05) that are also biologically relevant (i.e., contributing to the Dasatinib-resistance mechanisms). Including the above, the p53, Hippo, Neurotrophin, TNF, Thyroid hormone, and Sphingolipid signalling pathways were also found statistically significant (FDR-corrected *P*-value < 0.05). **(b)** Relationship of differential expression and pathway perturbation activity The Presence of potential drivers of the perturbation can be observed through either high perturbation score and/or significant expression changes. **(c)** Statistical significance of the negative total perturbation shows the potentially critical role of this pathway inhibition in dasatinib resistance mechanisms. **(d)** Pathview plots for the DEGs demonstrate their significant downstream effect contributing to dastatinib resistance mechanisms. These include downregulation of apoptosis (via CASP8), Increased inhibition of angiogenesis and metastasis, inhibition of IGF-I/mTOR pathway, etc.

**FIGURE 7.**
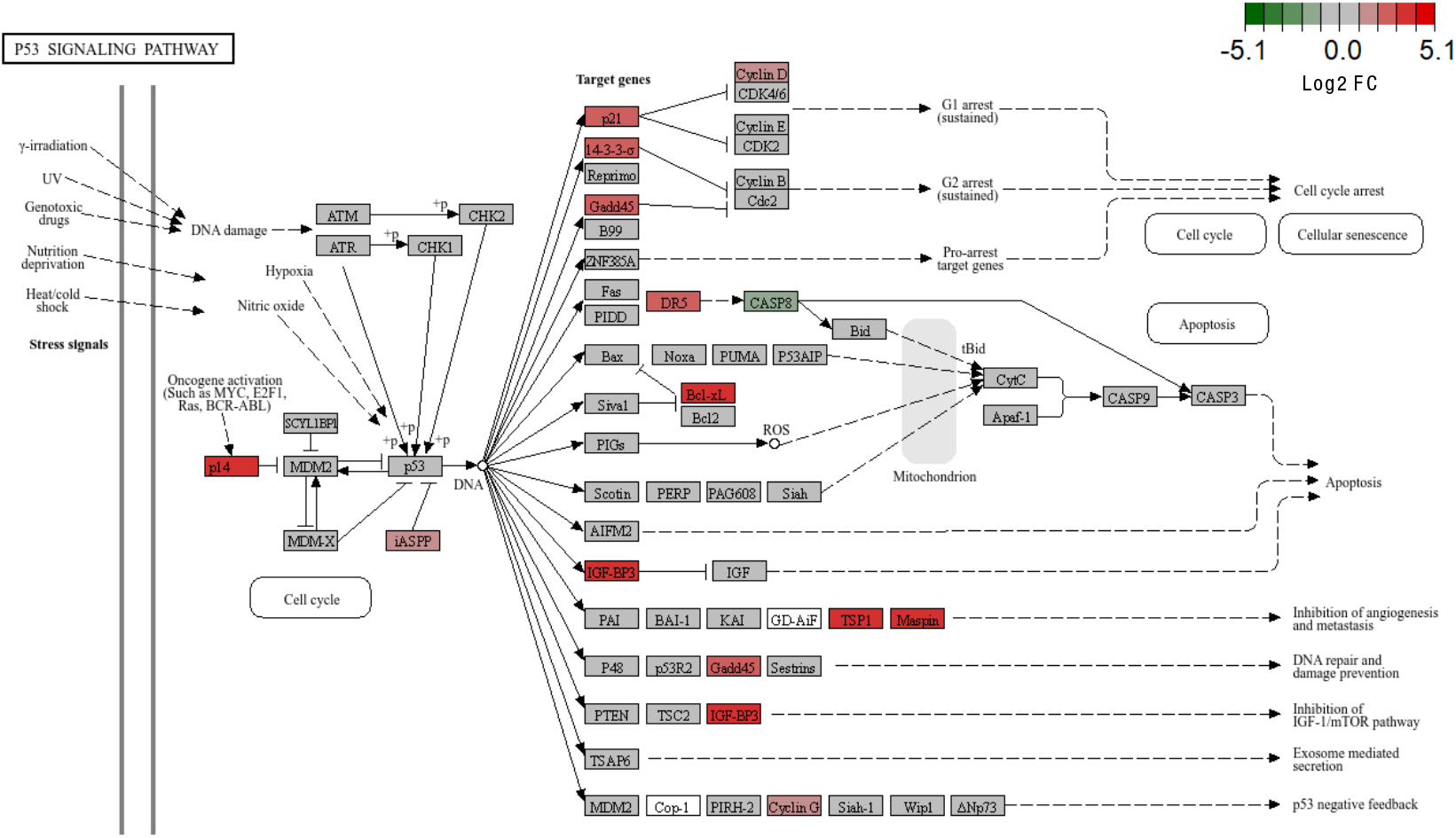
p53 Signalling pathway Perturbation. Pathview plot for the DEGs demonstrates their significant downstream effect contributing to dastatinib resistance mechanisms. These include downregulation of apoptosis (via CASP8), Increased inhibition of angiogenesis and metastasis, inhibition of IGF-I/mTOR pathway, etc .

### Validating with StringDB and Human Protein Atlas Databases

Identified resistant biomarkers [see Table 5] were further validated using existing knowledge based on StringDB, TCGA gene expression changes from the Human Protein Atlas, and PubMed. Using the StringDB web portal, 10 biomarkers were queried for potential connections among them - revealing an induced subnetwork of 10 nodes (confidence score threshold ≥ 0.4), where average node degree was observed as 1.8 and average local connected components = 0.633. This induced subnetwork also exhibited a PPI enrichment *P*-value = 0.015, indicating that the number of observed edges (9 edges) is significantly more than expected (4 edges) for a random gene set with the same size and same degree distribution [Figure 8a]. Interestingly, none of the StringDB connections found among these 10 resistant biomarkers were exhibiting physical protein-protein interactions, but only gene co-expression, co-mentioned in Pubmed abstracts, and homologs interactions in other species. However, functional enrichment of these resistant biomarkers with GO terms (Biological Process), Subcellular localizations (compartments) [see Figure 8b and Figure 8c], Reactome Pathways, WikiPathways, and Pubmed references [Supplementary Figure S1, Supplementary Figure S2, Supplementary Figure S3] revealed highly significant results with FDR-corrected *P*-value ≤ 0.05 [See Supplementary Table S1].

**FIGURE 8.**
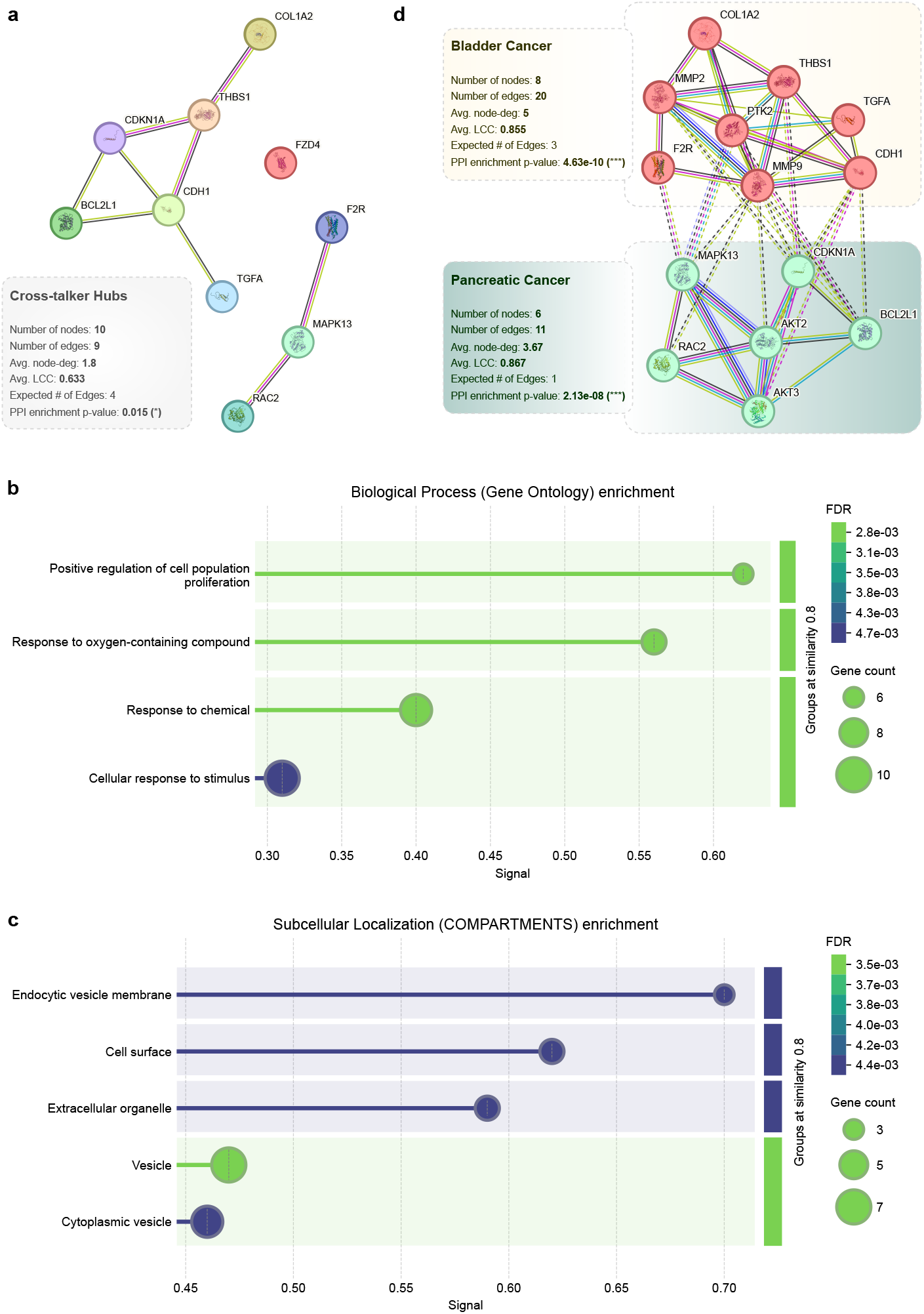
StringDB validation of 10 cross-talking biomarkers in Dasatinib resistance. **(a)** Protein-Protein interactions among cross-talking hubs (confidence score ≥ 0.4). Enrichment of 10 cross-talking biomarkers with **(b)** GO term enrichment (Biological Process) and **(c)** Subcellular localizations (compartments), indicating additional evidence and localized roles of these biomarkers in dasatinib resistance. **(d)** With augmented nodes and their incident connections from StringDB, an MCL clustering yielded two clusters, cluster-1: *Bladder cancer* (8 genes), and cluster-2: *Pancreatic cancer* (6 genes), each having statistically significant PPI enrichment scores.

Without physical connections, such significant functional relevance of these resistant biomarkers encourages us to observe beyond by enlarging the network (with a threshold of no more than 10 interactors) with the aim of identifying actual functional modules. Incidentally, by doing so, two clusters were identified by the MCL clustering algorithm [Figure 8d] - Bladder Cancer (additional genes: MMP2, MMP9, and PTK2) and Pancreatic Cancer (additional genes: AKT2 and AKT3), each having statistically significant PPI enrichment scores, 4.63 × 10^−10^ and 2.13 × 10^−08^, respectively.

A manual literature search with these findings revealed their connections with pancreatic cancer resistance mechanisms. While “Positive regulation of cell population proliferation” is highly relevant with pancreatic cancer resistance [20], its connection with the “Response to oxygen-containing compound” biological process has been multifaceted, which involves hypoxia adaptation [21], Reactive oxygen species (ROS) balance [22, 23], and stromal interactions [24]. Moreover, genes/protein alteration involved in “Endocytic vesicle membrane” compartments may influence drug trafficking and receptor internalization, with RCC1-mediated disruption of nucleocytoplasmic transport impairing metabolic pathways critical for chemotherapy efficacy [25]. The cell surface protein STK33, stabilized by the lncRNA MACC1-AS1, suppresses ferroptosis to evade chemotherapy-induced oxidative damage [26]. Cancer-associated fibroblasts (CAFs) mediate stromal-tumor crosstalk, transferring cytokines like IL-6 and oncogenic miRNAs that drive resistance [27].

Moreover, the resistant biomarkers were also evaluated with their potential coherence with gene (TCGA: The Cancer Genome Atlas) and protein expression (CPTAC: Clinical Proteomic Tumor Analysis Consortium) changes in cancer-vs-normal patients, collected from the Human Protein Atlas (HPA) database [13]. Among 10 biomarkers found in dasatinib resistance, only **RAC1, TGFA**, and **BCL2L1** were found prognostic in Pancreatic adenocarcinoma (PAAD) with statistical significance (*P*-values < 0.001) based on the TCGA gene expression data only [see Supplementary Figure S4].

Tumour-vs-normal comparison of relative protein expression data (nRPX) from CPTAC was conducted using the Wilcoxon test for all the resistant biomarkers, where 2 genes (FZD4 and TGFA) revealed their proteomics data as missing. Among the remaining 8 genes, all the genes demonstrated statistically significant (*P*-value < 0.001) differential *proteome* abundance in tumours-vs-normal, except F2R, CDKN1A and RAC2 [Figure 9]. Moreover, THBS1, COL1A2, and BCL2L1, were upregulated, whereas MAPK13 and CDH1 were found as down-regulated [Table 7]. Moreover, using RNA expression data from TCGA, only BCL2L1 and TGFA genes were found to be significantly prognostic [Figure 10].

**TABLE 7.**
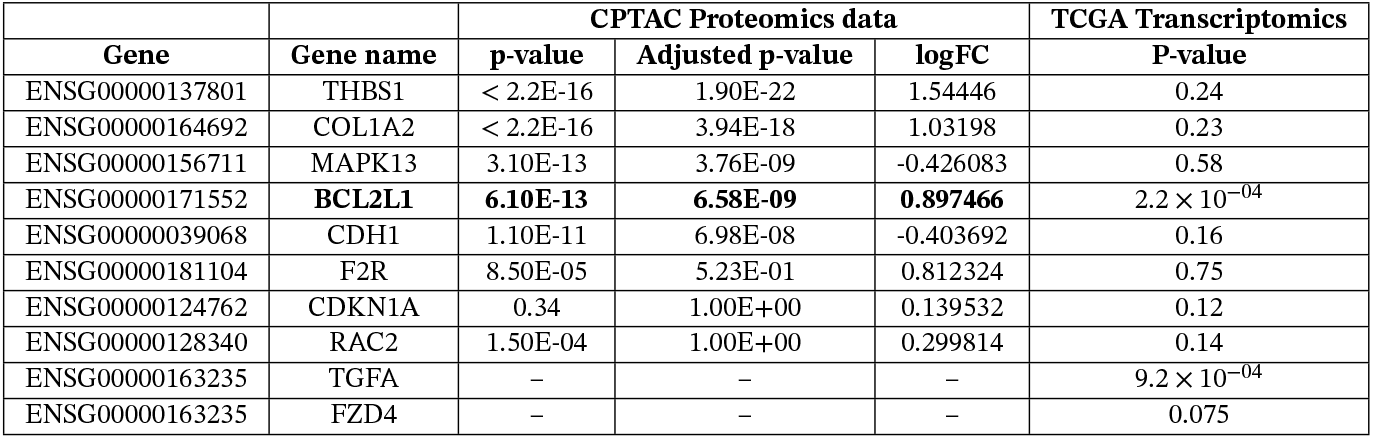
Validation of resistant biomarkers with CPTAC Proteomics and TCGA transcriptomics in PAAD.

| Gene | Gene name | CPTAC Proteomics data |  |  | TCGA Transcriptomics |
| --- | --- | --- | --- | --- | --- |
|  |  | p-value | Adjusted p-value | logFC | P-value |
| ENSG00000137801 | THBS1 | < 2.2E-16 | 1.90E-22 | 1.54446 | 0.24 |
| ENSG00000164692 | COL1A2 | < 2.2E-16 | 3.94E-18 | 1.03198 | 0.23 |
| ENSG00000156711 | MAPK13 | 3.10E-13 | 3.76E-09 | -0.426083 | 0.58 |
| ENSG00000171552 | **BCL2L1** | **6.10E-13** | **6.58E-09** | **0.897466** | $2.2 \times 10^{-04}$ |
| ENSG00000039068 | CDH1 | 1.10E-11 | 6.98E-08 | -0.403692 | 0.16 |
| ENSG00000181104 | F2R | 8.50E-05 | 5.23E-01 | 0.812324 | 0.75 |
| ENSG00000124762 | CDKN1A | 0.34 | 1.00E+00 | 0.139532 | 0.12 |
| ENSG00000128340 | RAC2 | 1.50E-04 | 1.00E+00 | 0.299814 | 0.14 |
| ENSG00000163235 | TGFA | – | – | – | $9.2 \times 10^{-04}$ |
| ENSG00000163235 | FZD4 | – | – | – | 0.075 |

**FIGURE 9.**
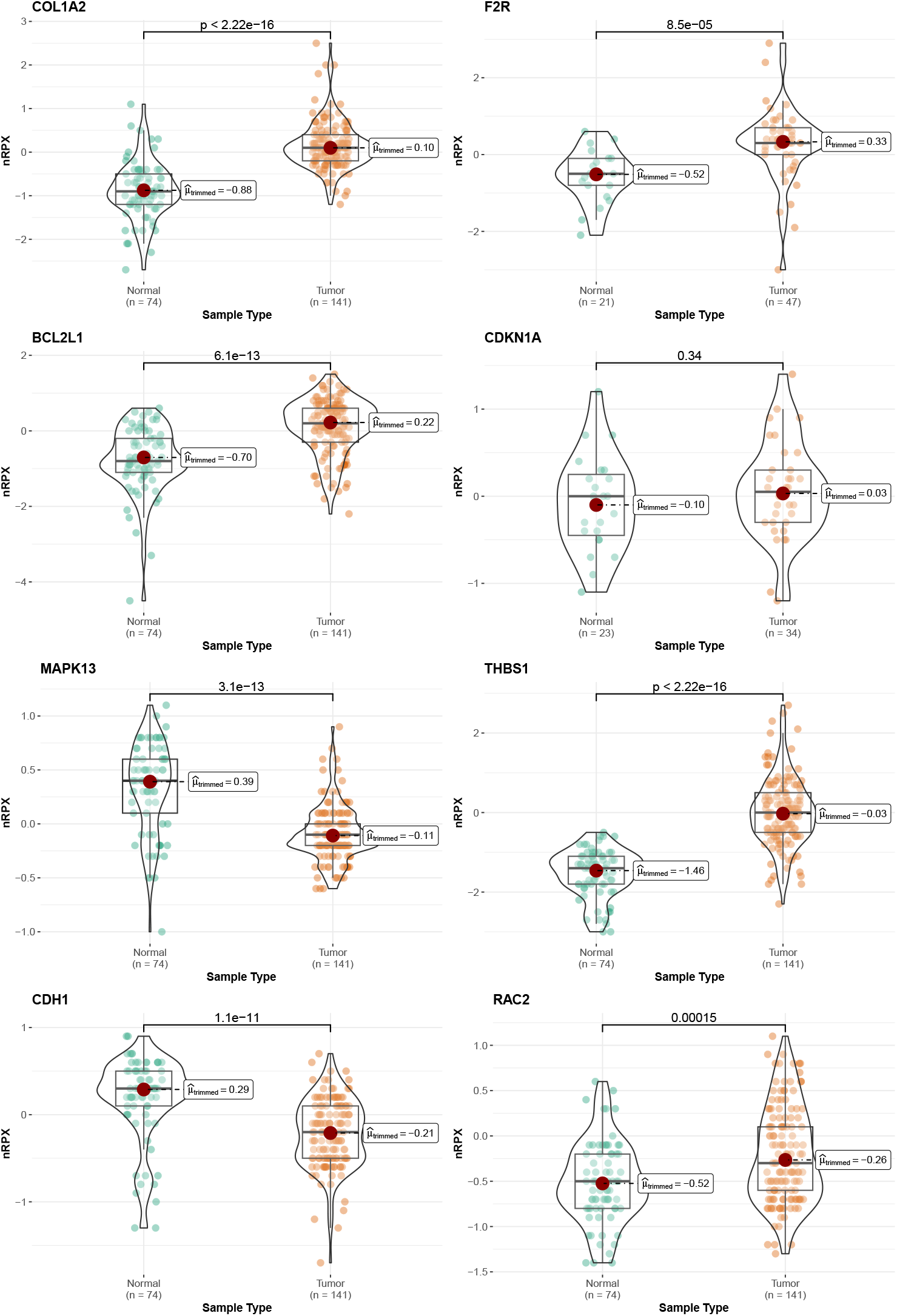
CPTAC relative proteomics validation of 8 or 10 resistant biomarkers. Most of the biomarkers (Except CDKN1A) were found to be significantly dysregulated (Wilcoxon test) in Pancreatic adenocarcinoma (PAAD) in Tumor-vs-Normal relative protein express (nRPX). Note 2 other genes, FZD4 and TGFA, weren’t found with any expression data.

**FIGURE 10.**
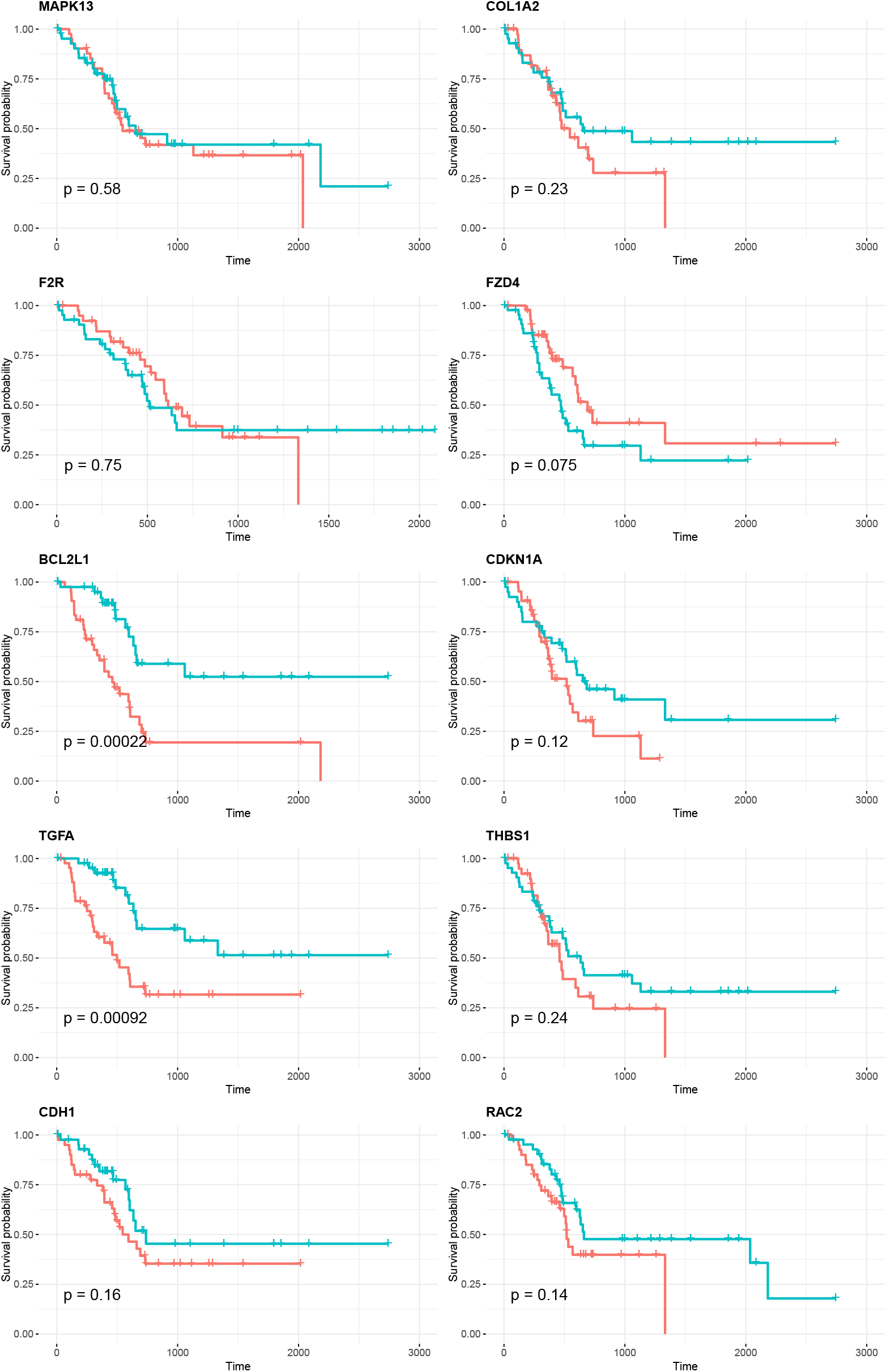
TCGA (RNA expression) validation 10 resistant biomarkers. Survival analysis was conducted using the Cox regression model.

## 4 Discussion & Limitations

Drug resistance is a significant threat in cancer treatment where resistant cells’ growths can not be controlled despite continuous intake of corresponding drugs. In this study, we analyzed an array-based transcriptomics dataset from dasatinib-sensitive and dasatinib-resistant cell lines to decipher the putative mechanistic details of dasatinib resistance in pancreatic cancer. Here, we conducted an integrated analysis of differential biomarker identification, functional term and pathway enrichment analysis to identify a tentative list of differential biomarkers and their associated pathways that revealed biological relevance - yielding *p53 signalling* and *FC epsilon RI signalling* pathways as the most significantly enriched. Gene set enrichment analysis (GSEA) identified terms that were predominantly related to pancreatic cancer, which suggests the biological relevance of DEGs in resistant-vs-sensitive conditions. Moreover, an ad-hoc and scale-free topology inducing inter-pathway cross-talk network analysis identified that these two pathways (i.e., *p53 signalling* and *FC epsilon RI signalling*) combinedly drive the overall perturbation by cross-talking with 87% (14 out of 16) all significantly enriched signalling pathways. During the signalling pathway impact analysis (SPIA), these two pathways were also found highly perturbed based on total perturbation accumulation and degree of dysregulation, where their Bonferroni-corrected *P*-values were 3.1 × 10^−5^, and 1.8 × 10^−2^, respectively [see Table 6].

P53 signalling plays crucial roles in mediating the normal growth of cells and utilizes its tumour-suppressing capability, however, re-programming of its signalling activities significantly facilitates dysregulation of normal cell functions, such as cell growth, apoptosis, DNA damage repair, and metabolism, which may infer resistance to dasatinib therapy [28, 29]. Regarding the FC epsilon RI (a high-affinity receptor of IgE) signalling, some literature suggests that the angiogenesis in pancreatic cancer can be induced by the inflammation caused within tumour micro-environment [30, 31]. However, the role of *FC epsilon RI signalling* activities in pancreatic cancer is poorly understood [32]. In our study, several genes in this pathway are differentially regulated that are responsible for pancreatic cancer metastasis and/or resistance to therapies including LYN [33], FYN [34, 35], RAC1 and RAC2 [36], MAPK13 [6].

Those two pathways managed to disseminate their perturbations globally via 13 cross-talking DEGs (*degree* ≥ 2), among which 5 of them showed higher degree of dysregulation (*abs*(*log*_2_*FC*) ≥ 2) in dasatinib resistance cell lines, including **THBS1, CDKN1A, BCL2L1, RAC2**, and **MAPK13**. THBS1 expression was found to be highly upregulated in pancreatic cancer cells, knockdown of which suppresses the pancreatic cancer progression in nude mice [37]. The cyclin-dependent kinase inhibitor p21 (CDKN1A) has multifaceted (dual-role) and context-dependent roles, including both tumour-suppressing and potential Pro-tumorigenic roles [38]. Alexandros *et al*. suggested that in p53 deficient micro-environment, p21 may play oncogenic roles depending on its subcellular localization [38]. Interestingly, in this study, we found the downregulation of TP53 (*log*2*FC* = −0.55) and BCL2 associated genes (BAD: *log*2*FC* = −0.67, BCLAF1: *log*2*FC* = −0.27, BAG2: *log*2*FC* = −0.23, BAG3: *log*2*FC* = −0.67, and BAG6: *log*2*FC* = −0.4) and the upregulation of MDM2 (*log*2*FC* = 0.37) in resistant-vs-sensitive cell lines, which indicate the overall downregulation of p53 signalling pathway. Moreover, the SPIA (Signalling Pathway Impact Analysis) results have independently identified the p53 signalling pathway as “*Inhibited*”, together which may suggest that p21 (CDKN1A) may acquire an oncogenic role in pancreatic cancer in resistance condition by inhibiting apoptosis and promoting abnormal cell growth [38]. Moreover, CDKN1A upregulation has been observed in cisplatin-pemetrexed chemoresistance in non-small cell lung cancer cells [39].

Over-expression of BCL2L1 was observed in pancreatic cancer patients [40], and it also induces resistance to several treatments, including gemcitabine, 5FU, and niraparib, in the pancreatic cancer cell line, AsPC-1 [41]. RAC2 promotes the tumour growth and metastasis [42], and activated RAC2 GTPase mediates Perineural invasion and metastasis in pancreatic cancer via MUC21 phosphorylation [43]. MAPK13 (p38 *δ*) is a member of the mitogen-activated protein kinase (MAPK) family, which works as a cross-talking signalling hub and plays crucial roles in cell proliferation, transcription regulation and differentiation. Godfrey *et al*. has reported that, due to epigenetic reprogramming of DNA hypermethylation, cells acquire resistance to MAPK signalling inhibitors in pancreatic cancer [6].

Identifying and analysing cross-talks among highly active signalling pathways can facilitate better mechanistic details underlying any disease context. To underscore the systemic nature of resistance mechanisms, here in this study, we modelled the signalling cross-talk network as a gene-pathway association, where the nodes are either DEGs or enriched signalling pathways where they belong and the edges were defined with the gene-pathway associations from known annotations, e.g., KEGG [Figure 4]. Therefore, cross-talking nodes were only those DEGs that were connecting at least two signalling pathways, and non-cross-talking DEGs were otherwise. For instance, p53 signallings cross-talk with pathways like ErbB, Wnt, and PI3K-Akt via key upregulated genes such as CDKN1A and BCL2L1 suggests its central role in orchestrating cellular responses to Dasatinib.

Note, Figure 4 shows only the pruned version of that network, where the DEG nodes that have a degree of connectivity as 1, were hidden as they weren’t effectively defined as cross-talking DEGs. Again, although this cross-talk network is constructed in an adhoc manner, we urge that it manifests biological significance since the gene-pathway membership edges indirectly map some meaningful biological connections (e.g., protein-protein interactions). Similar claim can be made based on the observation of its scale-free topology [Figure 5a-b] and the correlation of the degree of connectivity (i.e., node degree) and the degree of dysregulation (*abs*(*log*_2_*FC*) ≥ 2) for the cross-talking hubs [Figure 5c-d]. Since the cross-talking DEGs offer compensatory pathway activations contributing to the drug resistance phenomenon [8], detecting and targeting those are very crucial for novel therapeutics design. This study possesses several limitations. Firstly, we have used transcriptomics dataset only for discovering biomarkers in acquired dasatinib resistance (ADR), which may not fully reveal post-transcriptional or post-translational interactions among signaling proteins or molecules in that context. Secondly, proposed the cross-talk network model only relates the context with a static gene-pathway association, which may ignore dynamic scenario within tumor micro-environment. Moreover, cell-line derived transcriptomics datasets may also not capture the ADR mechanisms *in vivo* settings. To overcome these limitations, we aim to integrated multi-omics datasets, including proteomics and metabolomics data to capture more comprehensive details of ADR mechanims in pancreatic cancer. Moreover, time-course data, single-cell, or spatial transcriptomics data can also be integrated to reveal dynamic cell-cell or pathway-pathway communications for better cross-talk network modeling in ADR. Finally, additional *in vivo* based experimental validation may strengthen the findings of ADR biomarkers in pancreatic cancer.

## 5 Conclusions

In this study, we systematically addressed our core research questions by identifying dysregulated signalling molecules in dasatinib-resistant versus sensitive pancreatic cancer cell lines, interrogating their coordinated cross-talk across enriched pathways, and pinpointing dominant drivers of global perturbation. Through integrated differential expression profiling, enrichment analyses, signalling cross-talk network modelling, and topology-aware SPIA, we demonstrate that p53 and Fc-RI signalling axes act as complementary and dominant perturbation disseminators, collectively coordinating 87% of enriched pathway cross-talk. Scale-free topology analysis further prioritized ten cross-talking hub biomarkersmost notably BCL2L1, RAC2, MAPK13, THBS1, and CDKN1Awhose dysregulation correlates with network centrality and resistance-associated signalling cascades. Multi-layer validation using TCGA, CPTAC, and StringDB supports their biological and prognostic relevance. While the study is limited by reliance on in vitro transcriptomic models and pathway annotations that do not capture dynamic temporal rewiring or direct causal interactions, the convergence of statistical, topological, and biological evidence strengthens the robustness of our findings. Future work should incorporate single-cell and phosphoproteomic profiling, functional perturbation assays, and drug-combination validation to experimentally confirm these network vulnerabilities and enable rational design of next-generation therapeutic strategies targeting resistance dissemination rather than isolated molecular events.

## Data and code availability

- Gene expression data of dasatinib-resistant and dasatinib-sensitive pancreatic cancer cell lines are collected from Gene Expression Omnibus (GEO) under the accession number: GSE59357.
- All original code has been deposited at Zenodo (https://doi.org/10.5281/zenodo.14954417).

## Author Contributions

Conceptualization, A.A.; methodology, A.A.; investigation, A.A.; writing-original draft, A.A.; writing-review & editing, A.A. and M.A.M.; funding acquisition, M.A.M. and M.J.A.S.; resources, A.A.; supervision, M.A.M., and M.J.A.S.

## Acknowledgments

We thank Dr. Guangchuang (YuLabSMU) for his excellent R package: clusterprofiler, as some parts of the network visualization code in our work have been reproduced/inspired by that package.

## Supporting Information

Supplementary Figure S1, Supplementary Figure S2, Supplementary Figure S3, Supplementary Figure S4 and their legends in the supplementary files PDF

Supplementary Table S1. Full spreadsheet of differential expression of Resistant-VS-Sensitive pancreatic cancer cell lines.

