## Supplementary figures and images for "Global dissemination of FC-*ϵ* RI and p53 signalling perturbation contribute to Dasatinib resistance in Pancreatic Cancer Cell-lines"

### Supplementary Figure S1

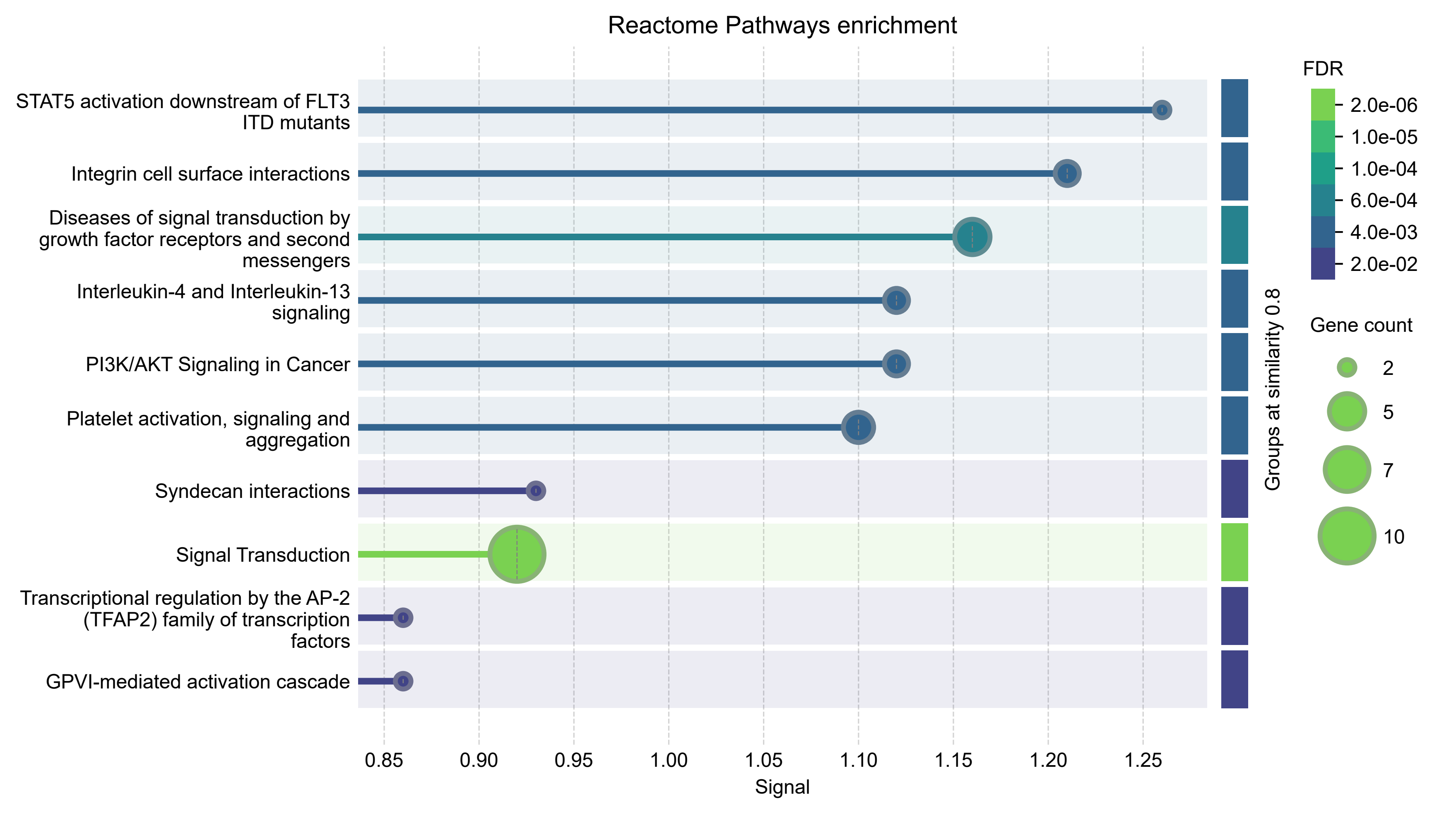

### Supplementary Figure S3

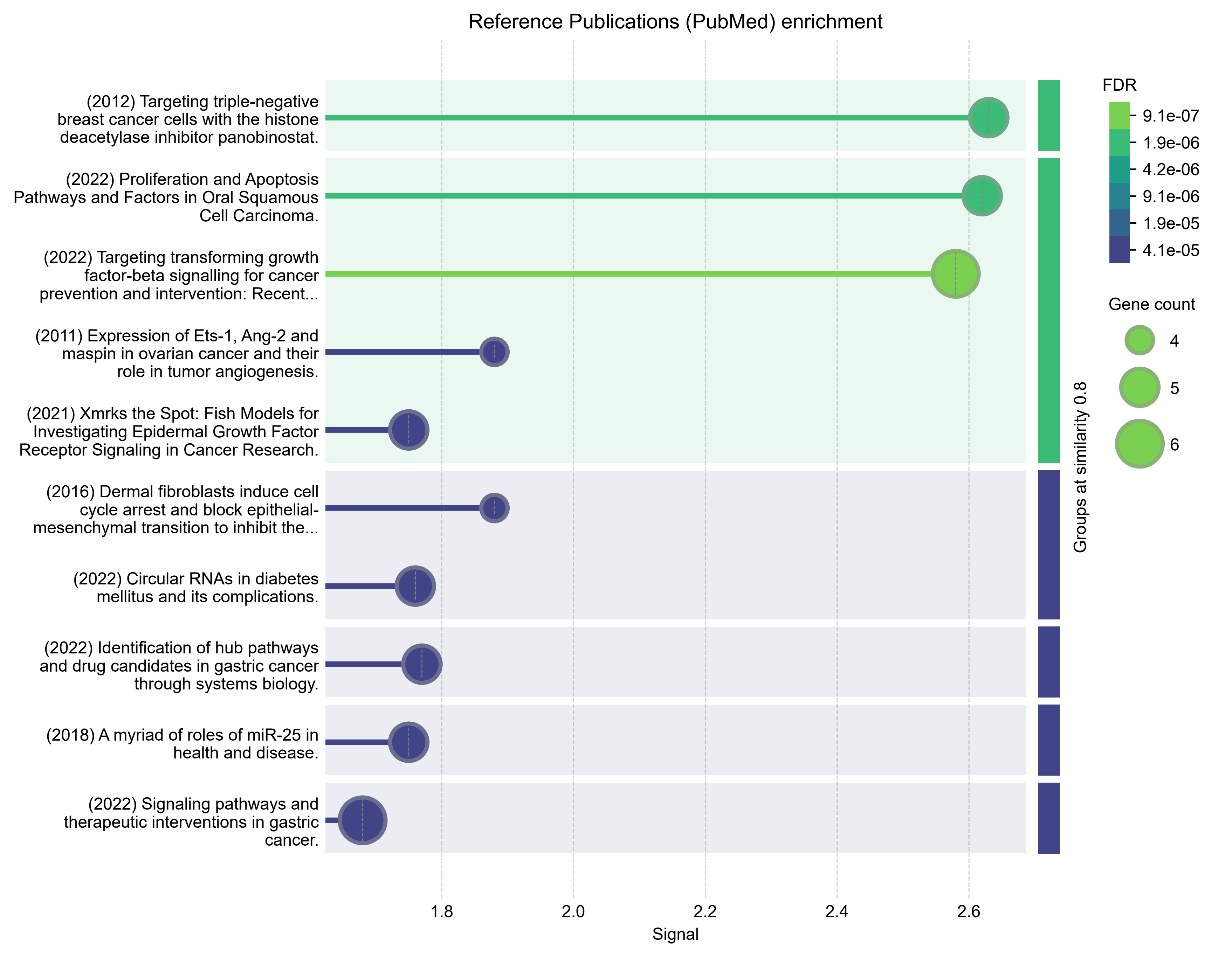

### Supplementary Figure S4

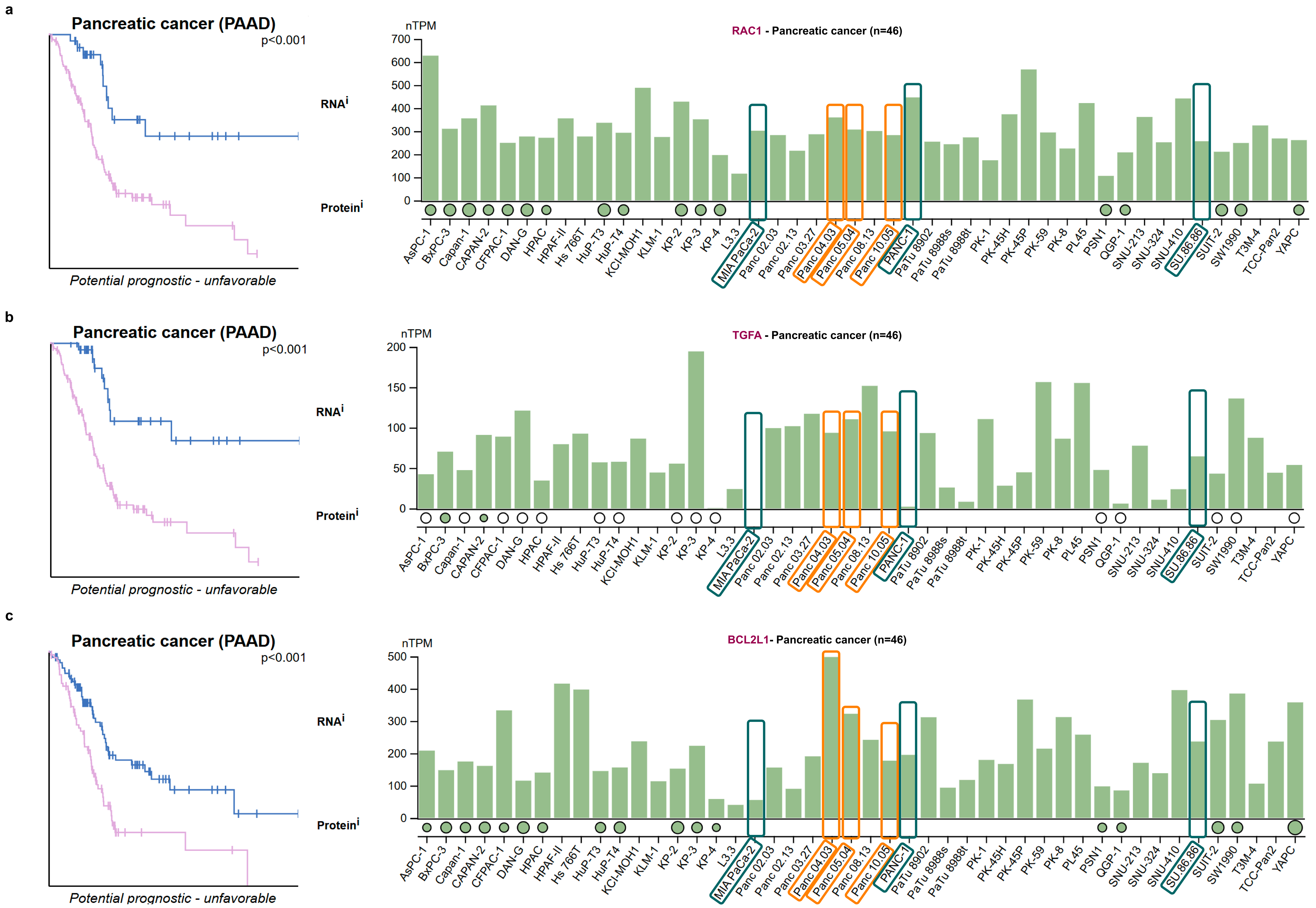
